# A novel naïve Bayes approach to identifying grooming behaviors in the force-plate actometric platform

**DOI:** 10.1101/2023.07.08.548198

**Authors:** Collin J Anderson, Roberto Cadeddu, Daria Nesterovich Anderson, Job A Huxford, Easton R VanLuik, Karen Odeh, Christopher Pittenger, Stefan M Pulst, Marco Bortolato

**Author notes:** **Corresponding authors: Collin J Anderson, PhD**, School of Medical Sciences, School of Biomedical Engineering, University of Sydney, Brain and Mind Centre, Room G505, 100 Mallett Street, Camperdown, NSW 2050, Australia, **Marco Bortolato, MD PhD**, Dept. of Pharmacology & Toxicology, College of Pharmacy, University of Utah, L.S. Skaggs Hall, Room 3916, 30 S 2000 E, Salt Lake City, UT 84112, USA.

## Abstract

**Background:** Self-grooming behavior in rodents serves as a valuable model for investigating stereotyped and perseverative responses. Most current grooming analyses primarily rely on video observation, which lacks standardization, efficiency, and quantitative information about force. To address these limitations, we developed an automated paradigm to analyze grooming using a force-plate actometer.

**New Method:** Grooming behavior is quantified by calculating ratios of relevant movement power spectral bands. These ratios are then input into a naïve Bayes classifier, trained with manual video observations. To validate the effectiveness of this method, we applied it to the behavioral analysis of the early-life striatal cholinergic interneuron depletion (CIN-d) mouse, a model of tic pathophysiology recently developed in our laboratory, which exhibits prolonged grooming responses to acute stressors. Behavioral monitoring was simultaneously conducted on the force-place actometer and by video recording.

**Results:** The naïve Bayes approach achieved 93.7% accurate classification and an area under the receiver operating characteristic curve of 0.894. We confirmed that male CIN-d mice displayed significantly longer grooming durations compared to controls. However, this elevation was not correlated with increases in grooming force. Notably, haloperidol, a benchmark therapy for tic disorders, reduced both grooming force and duration.

**Comparison with Existing Methods:** In contrast to observation-based approaches, our method affords rapid, unbiased, and automated assessment of grooming duration, frequency, and force.

**Conclusions:** Our novel approach enables fast and accurate automated detection of grooming behaviors. This method holds promise for high-throughput assessments of grooming stereotypies in animal models of tic disorders and other psychiatric conditions.

## Introduction

Most rodent species frequently engage in self-grooming to maintain coat cleanliness. This innate behavior is also often enacted in relation to other physiological processes, including thermoregulation, and key ethological domains, such as social communication, pheromone dispersion, and emotional regulation ^1, 2^. Environmental stress markedly increases self-grooming, possibly to counter arousal and reduce excitement ^3^. The syntax of self-grooming follows a cephalocaudal, stepwise sequence with a high level of complexity and microstructural organization ^4, 5^. Because of its highly consistent pattern, self-grooming is often studied to analyze the neural basis of complex stereotypies and as a phenomenological proxy to study perseverative hyperkinetic behaviors ^5, 6^. Emerging evidence points to the relevance of grooming stereotypies as a tic-associated manifestation based on face, construct, and predictive validity criteria ^6, 7^. For example, striatal activation has been shown to underlie tics ^8^ and grooming compulsions in mice ^9, 10^. Post-mortem studies in striatal samples from subjects with high-severity tics have shown a marked reduction in several families of striatal interneurons ^11, 12^. Building on this finding, targeted striatal depletion of cholinergic interneurons (CINs) in adult mice has been shown to increase the frequency of self-grooming responses to environmental stressors ^13^. We recently confirmed similar results in mice subjected to early-life CIN depletion (CIN-d) ^14^. Notably, we found that the exaggerated grooming responses in CIN-d mice were reduced by benchmark therapies for tic disorders, such as the dopamine D_2_ receptor antagonist haloperidol ^14^.

These findings underscore the importance of studying self-grooming responses in animal models of tic disorders. While previous work evaluating grooming behaviors has primarily relied upon manual observation of videos recorded from experiments, the growing need for high-throughput screening and greater reproducibility underscores the importance of developing new methods for automated grooming detection. Indeed, others have recently pursued video-based machine learning approaches to detect and distinguish facial and body grooming.^15^

Herein, we report on a simple automated approach for detecting excessive grooming stereotypies using CIN-d mice ^14^. In contrast to previous automation approaches, we used the force-plate actometer developed by Fowler and colleagues to track an rodent’s center of mass and enable automated, high-throughput quantification of behavior ^16^. We have previously used force plate actometery to quantify gait abnormalities in another mouse model of tic disorders ^17, 18^, and we have also modified force plate acometers to enable specific quantifications of gait ataxia, tremor, and other behaviors ^19–23^. In this work, we aim to make grooming analysis higher throughput, automated, and bias-free; further, while force-plate-based metrics may not distinguish facial vs. body grooming as easily as video-based approaches, we aimed to make our procedure capable of quantifying differences in grooming behaviors based on force, which may provide mechanistic insight. We trained a Bayesian classifier utilizing a combination of manually identified grooming times and spectral power across several bands obtained through continuous wavelet transformation of x, y, and force components acquired from the actometer. Using this classifier to characterize grooming vs. non-grooming behavior enabled high-speed, automated detection of grooming with high congruence with manual detection recorded from various experimental conditions and facilitated the quantification of grooming force, yielding further characterization of this stereotypy. Finally, this work adds to the face and predictive validation of the CIN-d mouse model of Tourette syndrome (TS) in an automated, inherently bias-free fashion. We anticipate that this approach will enable higher-throughput, rigorous, and reproducible therapeutic testing in rodent models of psychiatric conditions that feature modifications in grooming behaviors.

## Materials and methods

### Animals

Homozygous choline acetyltransferase (ChAT)-cre female mice (B6; 129S6-ChAT tm2(cre)Lowl/J) were obtained from the Jackson Laboratory (006410) and crossed with C57BL6/J male mice (The Jackson Laboratory, 000664; Bar Harbor, ME, USA). Genotypes were confirmed by PCR. Mice were maintained on an inverted 12-hour light / dark cycle, with lights off at 6:00 AM and lights on at 6:00 PM, at a controlled temperature of 22°C, and had ad-libitum access to food and water. All experimental procedures were approved by the institutional animal care and use committee of the University of Utah and complied with US Public Health Service guidelines on the care and use of laboratory animals. All efforts were made to minimize the number of animals used, as well as their suffering.

### CIN depletion

CIN depletion was induced in ChAT-cre hemizygous pups, as previously detailed^14^. Briefly, stereotaxic surgery to inject AAVs bilaterally was performed on postnatal day 4 under sterile conditions with standard procedures. Anesthesia was induced through hypothermia, and then pups were head fixed in a neonatal frame Stereotax (Stoelting; Wood Dale, IL, USA). The head was swabbed with 70% ethanol, and a 5 μL Hamilton Neuros syringe (Reno, NV, USA) was positioned above the bregma. The striatum was targeted using coordinates of 0.5 mm posterior, 1.6 mm lateral, and 2.8 mm ventral from bregma. The needle was advanced slowly to the injection site, and 0.25 μL of either A06 AAV or C06 control AAV^13^ was infused into the surrounding tissue hemisphere. The syringe was left in place for a minimum of 2 minutes after infusion and then slowly withdrawn. After surgical completion, the pup was placed on an isothermal warming pad. Once its body temperature and skin color normalized and it began to move, the pup was then returned to its mother, and maternal behavior was monitored as previously described ^24^.

### Drugs

Diphtheria toxin (DT) was purchased from Sigma-Aldrich (St. Louis, MO), prepared in a 0.9% NaCl solution, frozen until use, and administered via intraperitoneal injection at a dosage of 1 μg/kg at postnatal day 18. The D_2_ receptor (D_2_R) antagonist haloperidol was also obtained from Sigma-Aldrich, prepared in a 0.9% NaCl solution, and administered via intraperitoneal injection at a dosage of 0.5 mg/kg 20 minutes before motor data collection. The injection volume for all systemic administrations was 10 ml/kg.

### Behavioral testing

All grooming recordings were made in a dedicated behavioral testing room maintained at an illumination level of 100 lux, and all behavioral analyses were started 5 s after placing the animal in the chamber. All sessions took place within a force-plate actometer with a 42 × 42 cm inner chamber^16^, with both video-captured manual observation and automated detection of grooming occurring for all experiments. Data input was obtained from four load cells connected to a standard laptop using a National Instruments NI USB-6008 data acquisition card (Fig. 1A). Using this setup, we adapted previously written custom LabView code^19, 20, 22^ to capture voltage output from each cell at 1000 Hz, enabling center of mass tracking and applied force data collection at 1000 Hz (Fig. 1B); next, we adapted previously written custom Matlab code to automate the detection of grooming behaviors and quantify force applied during grooming, as described below. Three experiments were performed on two cohorts of mice: i) analysis of grooming in freely moving male and female CIN-d mice and controls; ii) analysis of grooming in CIN-d and control mice subjected to spatial confinement, using a clear polycarbonate cylinder (10-cm diameter and 30-cm height) as previously described ^25^; and iii) analysis of the effects of haloperidol on grooming behavior in CIN-d and control mice under spatial confinement. The freely moving and spatial confinement experiments included 7 male control, 11 male CIN-d, 7 female control, and 7 female CIN-d mice, each used in both experiments. One male and one female control and two female CIN-d mice escaped the spatial confinement and recordings were not completed; these mice are thus not included in analysis for the second experiment. The experiments with haloperidol were carried out using 12 CIN-d males/group, under spatial confinement conditions.

**Figure 1:**
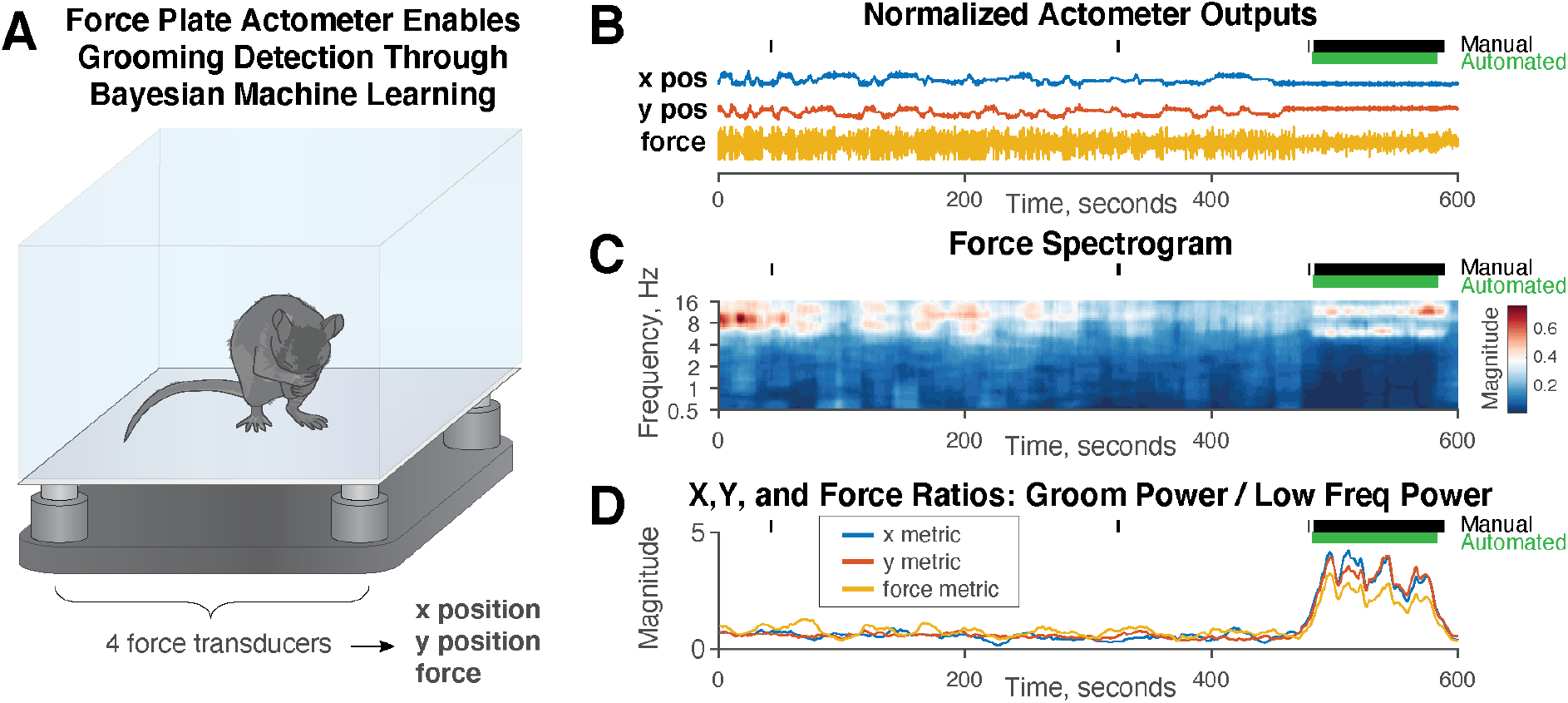
Method for automated detection of rodent grooming behaviors. **A.** Transduced force was captured from all four load cells of a force-plate actometer via a National Instruments DAQ card and fed into custom LabView code to track the center of mass and applied force at 1000 Hz before being analyzed in MATLAB. **B.** An example 10-minute recording shows traces of x and y positions as well as applied force, with both manually scored and naïve Bayes classifier-derived grooming identification above. Note that grooming was only identified during periods when the position remained steady, and overall fluctuations in force were reduced during grooming as compared to periods of ambulation. Example X, Y, and force plots are shown normalized with arbitrary units for the sake of simplicity. **C.** Based on Fourier analysis, it was clear that grooming occurs at 5-8 Hz. However, 5-8 Hz spectral power is often also strong during locomotion. However, <5 Hz power decreases when the rodent is not ambulating. Therefore, grooming corresponds with both increased 5-8 Hz power and decreased <5 Hz power. **D.** A ratio of 5-8 Hz power to 0.5-5 Hz power was computed for each of X, Y, and force and smoothed via a 10-second window convolution. Notably, this ratio was not identical across each orthogonal direction for all bouts of grooming, reflecting different orientations of both positioning within the chamber and grooming direction. All three ratios were then fed into a naïve Bayes classifier.

All 84 recordings (60 for the freely moving and spatial-confinement experiments and 24 for the haloperidol studies) were analyzed for grooming via both manual observation and the novel, actometer-based approach (see below). All sessions were video recorded and manually scored off-line for total grooming time over 10 minutes by a rater blind to experimental condition. Center of mass data recordings were simultaneously made using the force-plate actometer for all experiments. Behaviors considered to be grooming included complete and incomplete sequences of licking, scratching, and washing of the paws, head, body, and tail.

### Force-plate analytic procedures

Data were collected in National Instruments LabView and then imported into Matlab, as mentioned above. We adapted code previously first reported on by Anderson et al.^19^ and performed Fourier analyses on the center of mass tracking data. We had visually identified that grooming behaviors occurred at frequencies between 5 and 8 Hz; however, in comparing spectrograms to simultaneously capture video recordings of mice, we identified that 5-8 Hz power was often high not just during grooming but also during some periods of active movement (Fig. 1C). or even during occasional, brief periods of noisy signal. Therefore, given the strong 0.5-5 Hz power during both movement and saturating noise, we created ratios comparing 5-8 Hz power with 0.5-5 Hz power for movement in each of the X and Y directions, as well as for applied force (Fig. 1D). We applied a stratified random approach to select fifteen 10-minute actometer recordings. Using these grooming times as training data, we inputted all three directional Fourier ratios into a standard Matlab naïve Bayes model. We trained and tested a binary grooming classifier using a leave-one-out approach and then created a classifier based on all fifteen files. We applied this classifier across the full data set to identify groom times automatically. After grooming periods were auto-identified, we measured the average normalized force for each recording. The force applied to the load cells is primarily influenced by the mass of the animal, with greater gravitational force exhibited on the load cells by heavier animals. However, this force remains approximately constant during periods of rest, with greater variance from this baseline force generated by more forceful movement; thus, we computed normalized force based on how the scale of load cell output variability during grooming periods. We tested several methods for measuring the variability in force during auto-identified grooming periods with qualitatively similar results across each. Therefore, we used the simplest method in reported data, computing standard deviations across each automatically detected grooming period, then normalizing with 1.0 set as the average force applied by all male and female mice from the initial experiments in the open field, unrestricted conditions. Finally, we noted that some animals did not make detectable grooming behaviors during all recordings; as no grooming was detected, no grooming force can be extracted from these recordings, and these animals were removed from force analyses.

The naïve Bayes classifier was trained on 15 of the 84 experimental recordings selected via a stratified random sampling approach designed to include each experimental group reported in Figures 3-5 as well as short, medium, and long total groom durations as identified through manual observation. We tested the accuracy of the classifier on a leave-one-out approach 15 separate times, training the classifier on 14 files and testing it on the 15^th^. We created standard receiver operating characteristic curves for each of the 15 tests (Fig. 2A) and calculated the area under the curve for each. We calculated an average area under the curve across these values, as well as by micro, macro, and weighted average area under the curve methods. After retraining the classifier on all fifteen selected files, we performed a correlation analysis of manual observation vs naïve Bayes classification (Fig. 2B). All Matlab code and training model will be published with paper.

**Figure 2:**
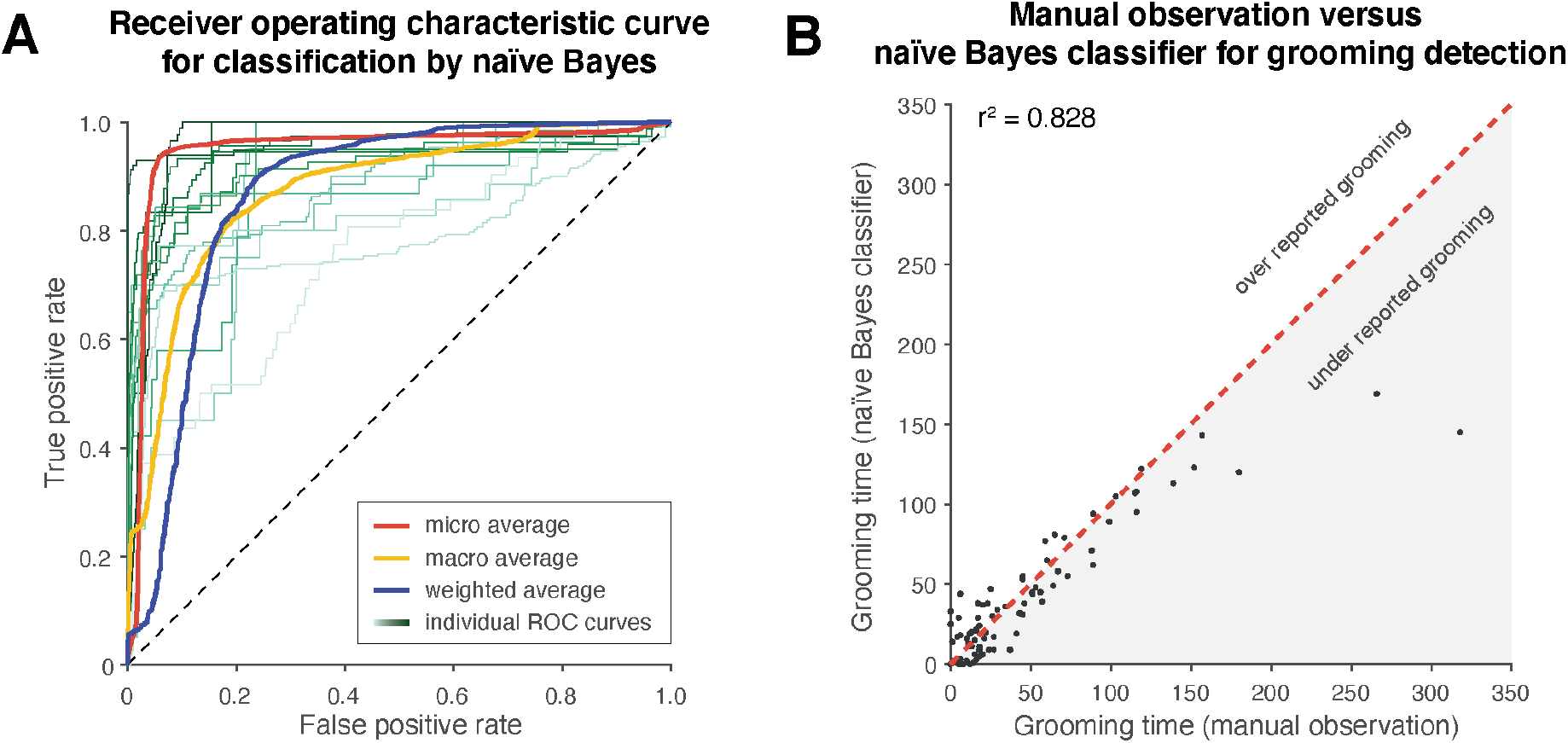
A naïve Bayes classifier automates the detection of grooming with high accuracy. **A.** 15 of the 84 10-minute experimental recordings were selected via random sampling stratified to include each experimental group reported in Figures 3-4, as well as short (bottom 10%), medium (middle 20%), and long (top 10%) total groom durations as identified through manual observation. We trained a naïve Bayes classifier on the 15 files and tested its accuracy on a leave-one-out approach. Thus, we tested the classifier 15 times by retraining it on 14 files and testing it on the 15^th^. Evaluating across all time points in the study, we found 93.7% agreement between manual grooming identification and the naïve Bayes classifier. We created a receiver operating characteristic curve for each of the 15 tests and achieved an average area under the curve of 0.894. Averages were recalculated via micro, macro, and weighted approaches, resulting in areas under the curve of 0.947, 0.873, and 0.867, respectively. This categorizes our classifier as either in the upper end of “excellent” or as “outstanding”^26^. **B.** Correlation analyses of manual observation vs. naïve Bayes classification yielded an r-value of 0.910 and an r^2-value of 0.828. Qualitatively, it appeared that the naïve Bayes classifier tended to miss very short duration grooming episodes (<3 seconds, as shown in Figure 1 B-D), likely due to the smoothing window applied to eliminate overclassification of noise as grooming; further, extreme outliers with grooming occurring 30+% of the time were not as well detected by the naïve Bayes classifier.

**Figure 3:**
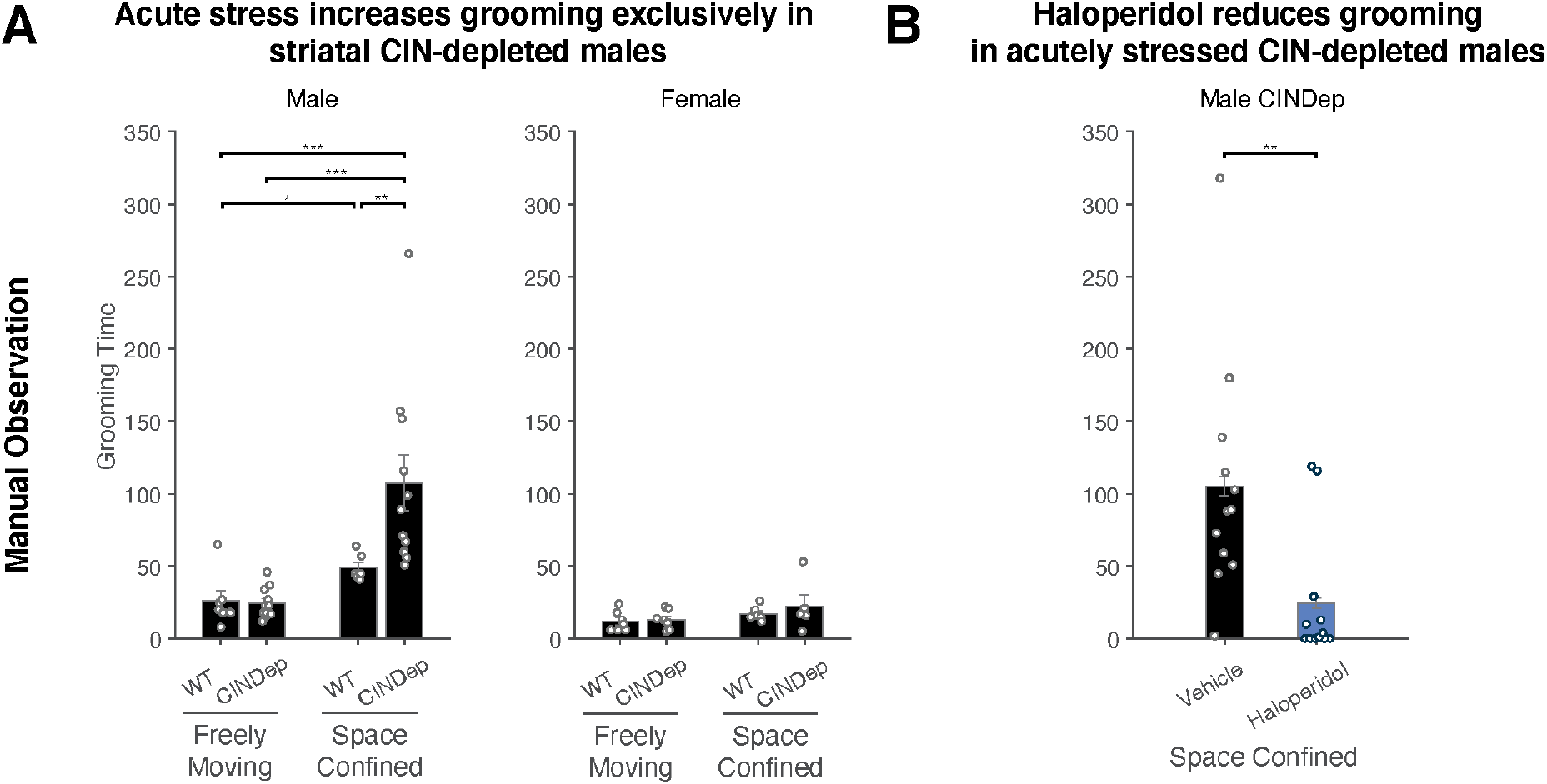
Striatal cholinergic interneuron depleted males, but not females, groom excessively in response to the acute stress of spatial confinement. **A.** Manual scoring of grooming from video by a blinded rater demonstrated that CIN-d males groomed excessively in response to the acute stress of spatial confinement. All other comparisons within sexes yielded insignificant differences. Error bars are only present for post-hoc tests; for ANOVA p-values, please refer to the text. **B.** The predictive validity of the striatal cholinergic interneuron depletion model is demonstrated: the D_2_ receptor antagonist haloperidol decreases excessive grooming made in response to spatial confinement.

### Statistical analyses

Statistical comparisons across groups were primarily performed via Two Way ANOVA; when significance was achieved, post-hoc comparisons were made using a Tukey’s honest significance test. In the case of a single comparison (Fig. 3B, 4B, 5B), a two-sample Student’s t-test was performed. Significance in correlation (Fig. 5C) was evaluated for using a Pearson’s correlation test. Significance thresholds were set at 0.05.

**Figure 4:**
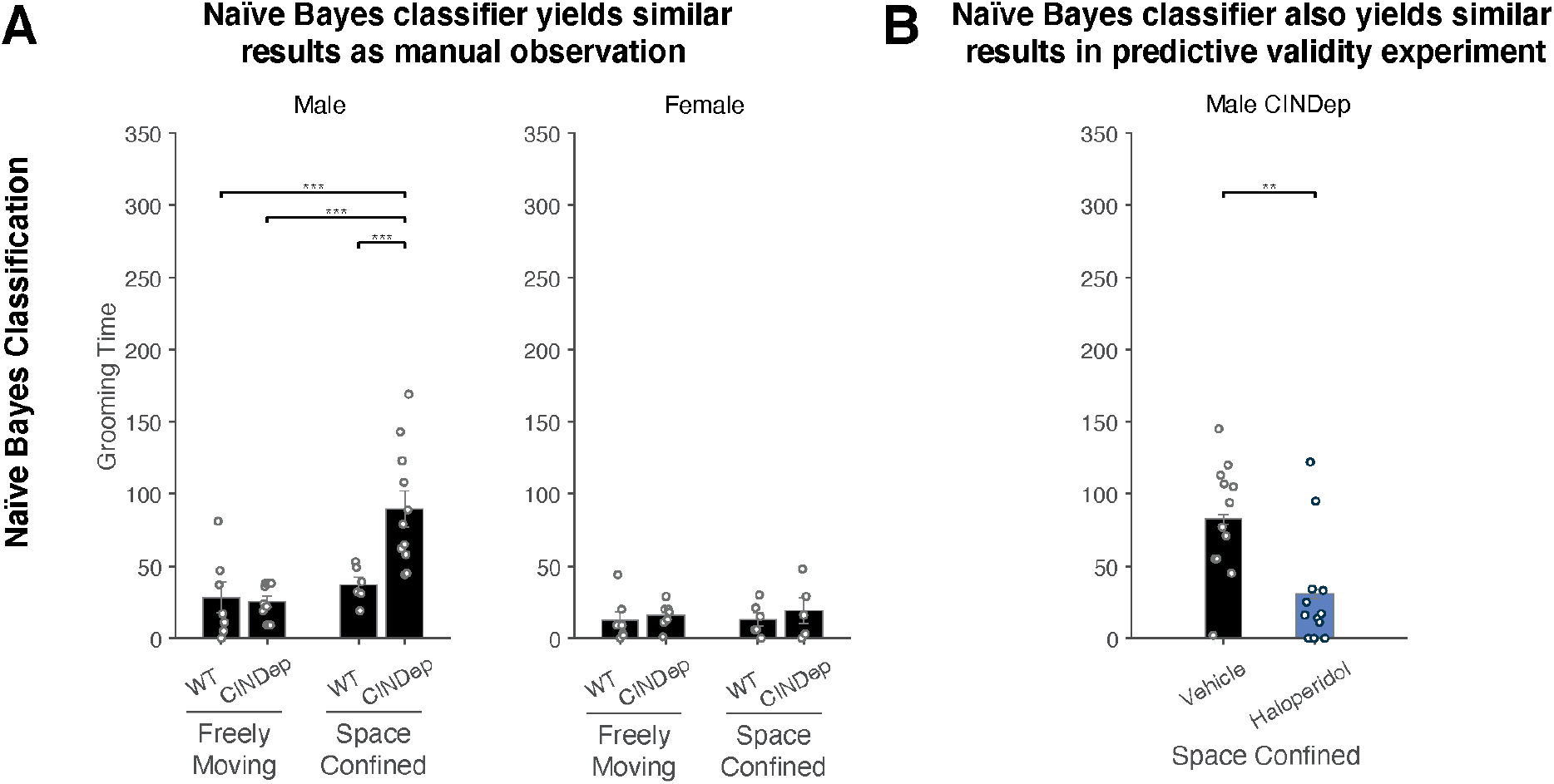
Naïve Bayes classification using force-plate data yields analogous results to manual observation. **A.** The experiment described in Figure 3 was reanalyzed using the naïve Bayes classification approach applied to force-plate data. Similar results were found, in that spatial confinement increased self-grooming in CIN-d males, but no other differences were found. Error bars are only present for post-hoc tests; for ANOVA p-values, please refer to the text**. B.** The D_2_ receptor antagonist haloperidol significantly decreased grooming time made in response to spatial confinement by CIN-d males, as was seen using manual scoring from video.

**Figure 5:**
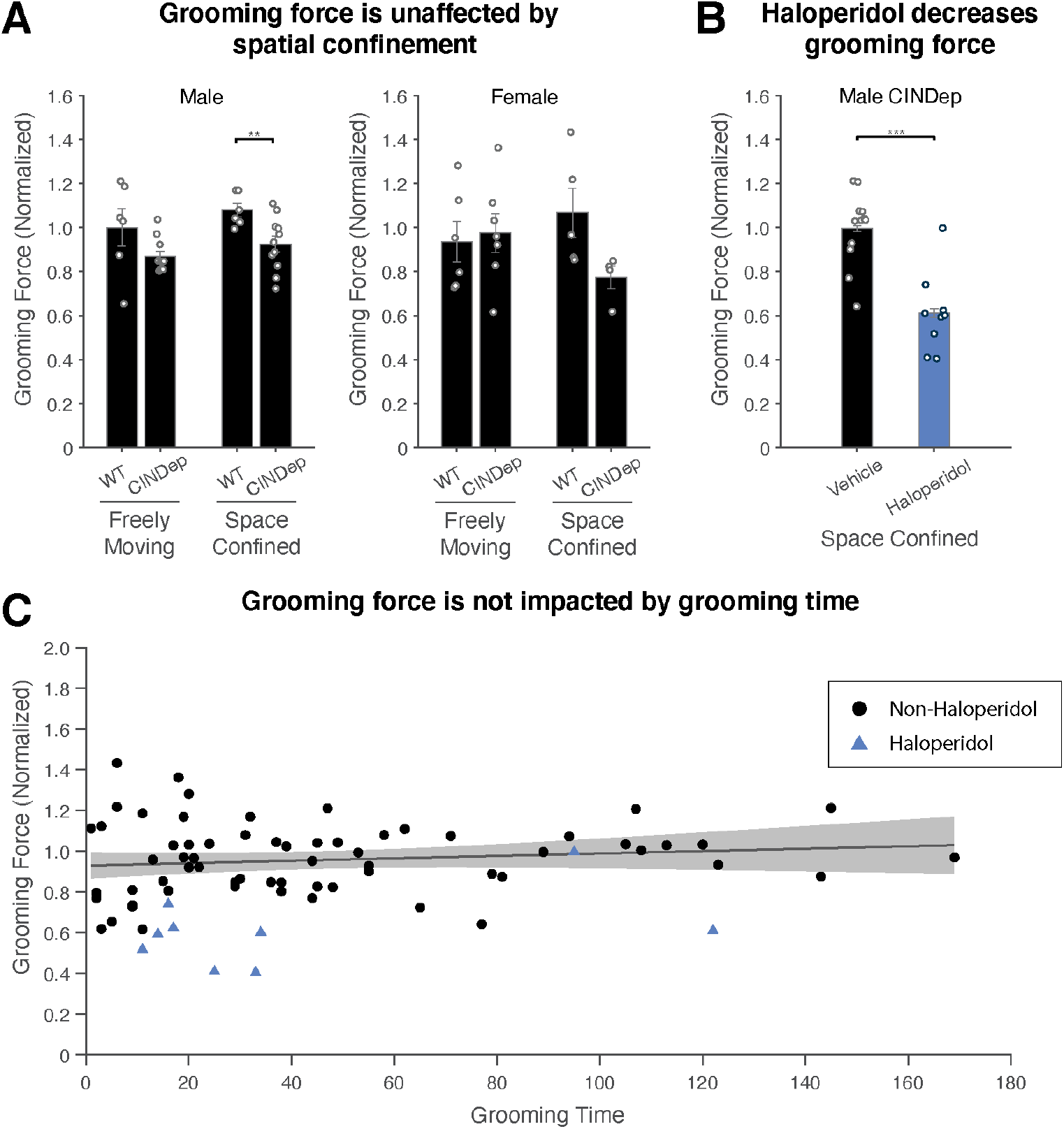
Excessive grooming is not tied to an increase in grooming force, although haloperidol reduces grooming force. **A.** We measured grooming force based on fluctuations in the net force applied to the load cells. The mean depression of the load cells scales with the mass of the animal, whereas fluctuations from this mean correspond to the forces applied during grooming episodes. Thus, we measured the variance of the net force applied to the load cells during grooming episodes to quantify the forcefulness of grooming. Grooming force was not significantly affected by acute stress through spatial confinement, although CIN depletion was found to reduce grooming force, with post-hoc tests showing a reduction in groom force specifically across genotypes during spatial confinement. **B.** Administration of 0.5 mg/kg of the D_2_ receptor antagonist haloperidol not only reduces grooming time (Figures 3-4), but also reduces grooming force. **C.** Pooling all experimental recordings from all studies, grooming time and grooming force are not significantly related. This conclusion is valid with or without the inclusion of data from haloperidol-treated mice.

## Results

### Manual video-based analysis of spontaneous and stress-induced grooming behavior

We quantified grooming in intact mice and mice in which striatal cholinergic interneurons (CINs) were experimentally depleted early in postnatal development (CIN-d). Quantification was first performed by manual scoring from video, blind to experimental condition. Self-grooming was evaluated in each mouse under freely moving conditions and under spatial confinement within a Plexiglas cylinder (Fig. 3A). Both spatial confinement (F(1,31)=23.26, p<0.001, η^2^=0.43) and CIN-d status (F(1,31)=4.95, p=0.034, η^2^=0.14) led to increased grooming in male mice, with a significant interaction (F(1,31)=4.28, p=0.047, η^2^=0.12). Post-hoc comparisons showed that spatial confinement elicited a mild yet significant elevation in grooming behavior in intact (p=0.017) and CIN-d mice (p=0.00154). The effect was more pronounced in CIN-d males (CIN-d vs control males under spatial confinement, p=0.013). Conversely, freely moving CIN-d and control mice exhibited comparable grooming behavior. In female mice, in a separate Two Way ANOVA, confinement stress mildly elevated grooming behaviors (F(1,22)=4.43, p=0.047, η^2^=0.07), but CIN-d status did not yield a difference, and there was no significant interaction. Post-hoc tests did not reveal significant differences across groups. In a separate cohort, CIN-d male mice were tested under spatial confinement conditions with an injection of haloperidol (0.5 mg/kg) or its vehicle (Fig. 3B). As expected, haloperidol significantly decreased grooming duration (t(22)=3.02, p=0.004).

### Validation of the naïve Bayes approach

We trained the naïve Bayes classifier to identify manually identified grooming from force-plate data through fifteen 10-minute experiments selected by stratified random sampling and then evaluated the classifier using a leave-one-out approach. The automated naïve Bayesian approach for detecting grooming yielded 93.7% agreement with manual grooming identification across all time points. We next evaluated the naïve Bayes classifier across these experiments on a leave-one-out approach, as described above. Creating receiver operating characteristic (ROC) curves for each file, we achieved an average area under the curve (AUC) of 0.894.

Recalculating averages using micro, macro, and weighted approaches yielded 0.947, 0.873, and 0.867, respectively (Fig. 2A). Based on the previous characterization of ROC curves for machine learning models, this rates our classifier as “excellent” or “outstanding”^26^. Comparing all 84 experiments recorded, we found that manual observation and naïve Bayes classification were very highly correlated, with an r-value of 0.910 and an r^2^-value of 0.828 (Fig. 2B). On average, the naïve Bayes classifier under-detected grooming by ∼11.89%; we primarily noted discrepancies in the context of very short (<3 s) bouts of grooming (as shown in the example in Fig. 1 B-D) and three recordings during which the animal groomed for more than 30% of the total time. Notably, the removal of the three most extreme length duration manually identified grooming sessions increased the r-value of 0.933 and the r^2^-value to 0.871, and reduce the average under-classification to only 4.21%.

### Automated actometer-based analysis of spontaneous and stress-induced grooming behavior

Having validated the naïve Bayesian classifier, we next re-quantified grooming in all experiments, with grooming times identified by the classifier (Fig. 4A). Under baseline conditions, as before, both spatial confinement (F(1,31)=21.88, p<0.001, η^2^=0.41) and CIN-d status (F(1,31)=6.51, p=0.0158, η^2^=0.17) led to increased grooming in male mice, with a significant interaction (F(1,31)=6.99, p=0.0127, η^2^=.18). In a separate Two Way ANOVA, no factors significantly changed grooming behaviors in female mice. In post-hoc tests, we found that control male mice did not exhibit a significant difference in self-grooming when spatially confined. However, CIN-d male mice engaged in longer grooming duration when spatially confined (p<0.001). Under freely moving conditions, no difference was found based on CIN-d status, while under spatial confinement conditions, CIN-d male mice exhibited more grooming than control mice (p=0.00192). Given that no factor yielded a significant difference in female mice on ANOVA, no post-hoc comparisons were made.

Finally, in the predictive validity experiment, we once again found results in agreement with the manual video-based analysis (Fig. 4B): haloperidol was found to significantly decrease grooming times (t(22)=3.26, p=0.00181).

### Automated actometer-based analysis of applied force during grooming

Having validated the naïve Bayes classifier applied to force-plate data as an automated method for detecting grooming behaviors, we proceeded to analyze grooming force across all experiments. In male mice, we found that CIN-d mice exhibiting a decrease in grooming force (F(1,30)=10.55, p=0.00287, η^2^=0.26). Stress did not significantly modify grooming force, and there was no significant interaction. No factors were significant in female mice. In post-hoc tests in male mice, we found no difference in any comparisons apart from a CIN-d status difference in spatially confined animals (p=0.00570). Importantly, spatial confinement itself did not modulate grooming force in CIN-d mice. There was no significant relationship between grooming time and grooming force (Fig. 5C).

## Discussion

Animal models are powerful research tools to advance our understanding of the pathophysiology of tic disorders. Several models have been developed based on observations in patients ^13, 14, 27^. The development of automated systems for assessing tic-like behaviors is critical to enable high-throughput screening of the behavioral output of these models. Here, we used a well-validated mouse model of tic pathophysiology and studied its effects on grooming using a force-plate actometer. This apparatus has been extensively used to quantify other motor behaviors in animal models of other neurological diseases, including ataxia, Parkinson’s disease, and Huntington’s disease. Indeed, we previously used this apparatus to study gait abnormalities in another animal model of tic disorders, the D1CT-7 transgenic mouse, and documented fine locomotor alterations in this mouse ^18^. Studies using the actometric platform on different animal models have shown that this apparatus can analyze several parameters of locomotor manifestations, including distance traveled, rotations, low mobility bouts, rearing, jumping, ataxia, and tremor.^28^ We have previously used force-plate actometers to quantify locomotor behaviors ^17, 29^, tremor ^19–21^, and ataxia ^19, 22^, and we have used actometer-derived data to support deep learning-based quantification of video-based tremor analyses ^20^. The work reported herein expands the body of reported capabilities of actometric data in animal models of tic disorders as applied to grooming stereotypies.

Using a well-validated mouse model of tic pathophysiology, we tested it in the force-plate actometer and found that a naïve Bayes classifier with inputs consisting of simple ratios of Fourier power in three orthogonal directions can rapidly and accurately detect self-grooming stereotypies. This approach allowed us to replicate our recent findings that male CIN-d mice display increased grooming in response to spatial confinement stress ^14^. Specifically, in our recent report, we documented that CIN-d males, but not females, exhibited higher grooming and other tic-related abnormal behaviors, which were significantly reduced by haloperidol. The automated analyses presented in this study validated all those results and showed that the increases in grooming exhibited by CIN-d males are not tied to increases in grooming force. Furthermore, haloperidol reduced both grooming duration and force, in line with well-established evidence on the therapeutic effects and motoric impairments induced by D_2_ receptor antagonists.

These experiments exemplify the use of a novel machine-learning approach to classifying grooming behavior. Compared to manual video-based analyses, this naïve Bayes approach to grooming quantification is inherently less biased and much faster. A full day of experiments can be analyzed in a fully automated fashion in approximately 15 min on a standard laptop. This will enable higher throughput and more rigorous testing of therapeutic strategies for tic disorders. Beyond therapeutic testing, this method provides utility for characterizing other novel rodent models of tic disorders. Finally, the inclusion of a force metric enables investigators to evaluate pathological and therapeutic mechanisms better. For example, this study showed that stress-induced increased grooming does not likewise increase in force from spontaneous grooming behaviors in control and CIN-d mice. However, haloperidol reduced not only grooming stereotypies but also grooming force, which may be relevant to its mechanism of action. Applying similar approaches reduces bias and increases the rigor we evaluate models. Machine learning approaches have generated automated approaches to the analysis of rodent models in a variety of contexts, including pose estimation for behavioral and motor quantification ^30^, ultrasonic vocalizations ^31^, tremor ^20^, and social behavior ^32^. Increased automation of behavioral analyses will increase rigor and substantially speed up computations, enabling many higher-throughput analyses of potential therapeutics for tic disorders.

Having the ability to quantify force of grooming in addition to grooming time is both interesting and useful. The relationship between the incidence, duration, and vigor of abnormal movements varies across hyperkinetic disorders. In some cases, such as with blepharospasm versus a typical eyeblink, the hyperkinetic movements can occur with a great deal more force^33^. Our findings here support the idea that tics and tic-like manifestations generated by striatal cholinergic pathology are not more forceful, but rather, simply more frequent and inappropriately timed. This informs upon the role of cholinergic interneurons and suggests that such tics and tic-like manifestations are not dystonic in nature.

A number of limitations should be acknowledged. First, machine-learning classifiers often yield accurate results across a variety of data types, but they are imperfect. In this case, our classifier had a tendency to miss very short grooming episodes, particularly those under 2 s, as illustrated by the example data in Fig.1 B-D. This likely occurs as a result of smoothing the data with a short running average window to avoid noise and overclassification as grooming. Second, we note that automated and visual analyses did not always match precisely on start and stop times of bouts of grooming, with disagreement by one or even several seconds relatively frequently. Third, we note that the three most extreme outliers in terms of very long groom durations – 2.4+ standard deviations above the mean, or the top ∼1% based on mean and standard deviation – were under-classified by our algorithm. This may be due to either the classifier not being trained on any extreme outliers or based on simply based on average values in the Fourier ratio input being so high across the recordings. Finally, while this method can quantify force metrics that would not be possible to quantify with video, it is worth noting that our method does not distinguish between different grooming components. Much work has been done in the context of grooming microstructure ^34^, and our method is not currently equipped to identify changes within the syntactic chain pattern or syntactic vs. non-syntactic grooming. This is a direction of interest for future work.

Even with these limitations, our results showed that using a naïve Bayes classifier with inputs pertaining to center-of-mass data collected on a force-plate actometer enabled the automated quantification of spontaneous grooming behaviors. This work validates our recent findings reported in the context of the striatal cholinergic interneuron depletion model of tic pathophysiology ^14^, and adds to the rigor with which we evaluate such models.

## Acknowledgments

This study was supported by the NIH grant R21 NS108722 (to MB and CP) and NIH grant R35 NS127253 (to SMP).

